# Tracing of the developmental origin of the caudal fin muscle in zebrafish

**DOI:** 10.1101/2024.12.13.628295

**Authors:** Kinya G Ota, Gembu Abe

## Abstract

Teleost species possess complex caudal musculoskeletal systems. While mid-trunk muscles exhibit simple segmental patterns, several caudal skeletal muscles display intricate orientations in their muscle fibers. Due to this distinctive morphology, both early and recent researchers have studied the structure and development of the caudal musculoskeletal system. However, the early developmental origin of the cell populations within the caudal muscle system remains largely unknown. In this study, we performed lineage tracing of caudal muscle primordia in zebrafish using a transgenic line expressing EGFP in somite derivatives following tamoxifen induction. This approach allowed us to observe the specific cell populations that contribute to caudal muscle tissue formation at the early larval stage. By monitoring the growth of these labeled cells from the early larval stage, we identified the origins of muscle fibers in caudal fin muscles unique to teleosts, such as the adductor caudalis and flexor caudalis. Our findings provide descriptions that aid in understanding how fish-specialized caudal muscle structures were formed through the modification of developmental processes during evolution.

## BACKGROUND

The post-cranial muscle system in the teleost species is highly differentiated in the caudal region. While the trunk muscle system consists of metamerically arranged muscle tissues, the arrangement and orientation of these tissues are significantly altered at the caudal levels (Flammang and Lauder, 2009; Greene and Greene, 1914; Lauder, 1989; Schneider and Sulner, 2006; Siomava and Diogo, 2018; Winterbottom, 1973). Although several caudal muscles are homologous with trunk muscles, certain caudal muscles—including the flexor caudalis ventralis, adductor caudalis, and intervertebral muscles—show substantial changes in their fiber orientations. Because of its unique morphology, physiological and anatomical studies have been conducted to understand these muscles’ functions and morphological adaptations (Flammang and Lauder, 2008, 2009; Lauder, 1989; Siomava and Diogo, 2018).

Despite numerous studies on skeletal development in the caudal and embryonic development of trunk muscles, detailed observations of the developmental origins of caudal muscle groups remain limited (Blagden et al., 1997; Devoto et al., 1996; Du et al., 1997; Faustino and Power, 1998; Holley, 2007; Kumar et al., 2024; Lleras Forero et al., 2018; Nguyen-Chi et al., 2012). To our knowledge, only a few studies—such as Siomava and colleagues (2018), who used phalloidin staining to observe caudal muscle development in zebrafish, and Ishikawa (1990), on medaka—have investigated caudal muscle development. However, these studies focused on relatively late-stage larvae, where muscle fiber differentiation was already well underway, leaving the early origins of caudal muscles largely unexamined.

To address this, we performed genetic mosaic experiments using a transgenic zebrafish line with somite-derived cells randomly labeled (Fig. 1). We generated this line by employing an enhancer region from the medaka *mesp-b* gene (Terasaki et al., 2006), which is well-known for its expression in the presomitic mesoderm and essential regulatory elements, due to its simpler genome structure compared to zebrafish (Terasaki et al., 2006; Yabe et al., 2016). Using this enhancer, we constructed the Tg(*Olmespa-ERT2-Cre-ERT2*) (*mespCre*) transgene and introduced it into zebrafish. By crossing these transgenics with a Tg(*P067Olactb*) (*actbRG*) line (Yoshinari et al., 2012), we created the “mespRG zebrafish” (Fig. 1). In this line, the *mespCre* transgene drives the expression of a fusion Cre recombinase-ERT2 protein, which, upon tamoxifen induction, transitions into the nucleus to excise the *loxP* sites in the *actbRG* transgene, switching the reporter fluorescent from DsRed2 to EGFP (Fig. 1A).

**Fig 1.**
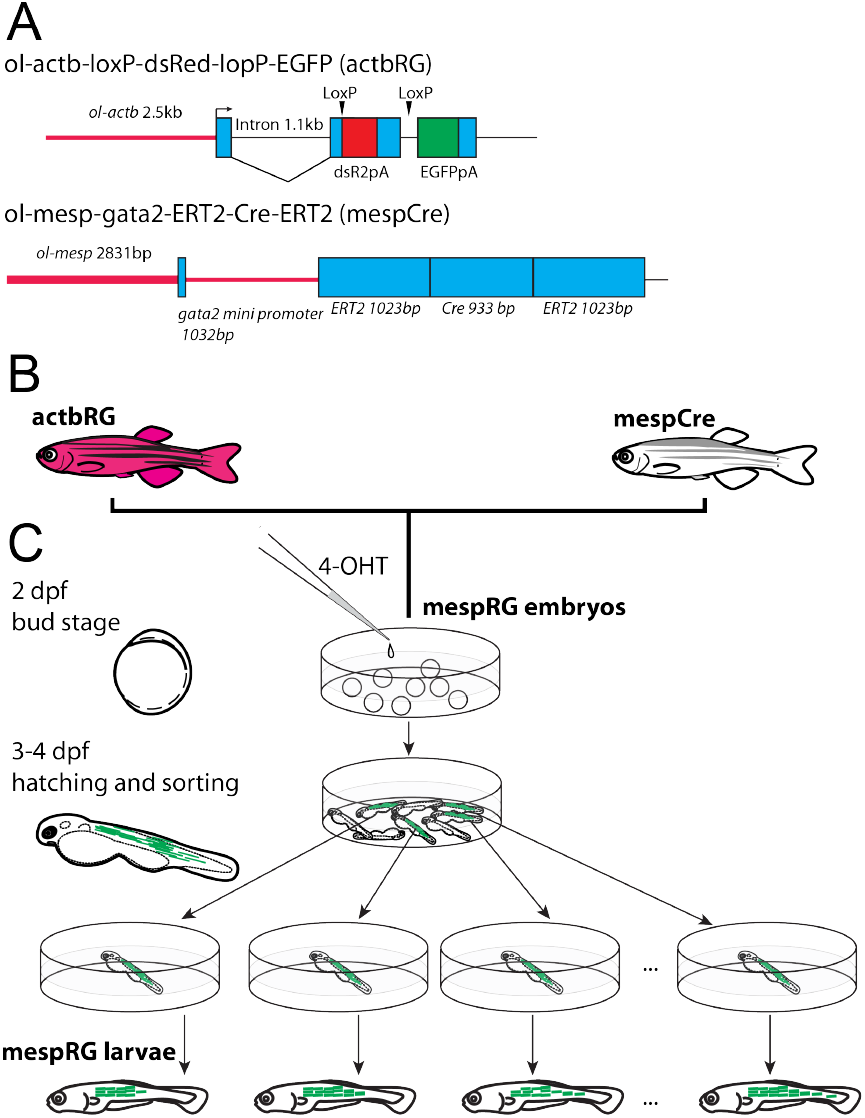
Genetic mosaic analyses of somite derivatives. A: Schematic representation of *P067Olactb:loxP-dsR2-loxP-EGFP* and *Olmespa-ERT-CreERt2* system. B. Parents zebrafish. C. mespRG population. Fish strains are indicated in bold letters (see Materials and Methods for details).

By labeling these precursor cells in the mespRG zebrafish, we tracked the appearance and migration patterns of muscle precursor cells in the caudal region. Our results offer insights into how the developmental processes of caudal muscles have been modified within teleost lineages.

## MATERIALS AND METHODS

### DNA construct

The *mespCre* construct was generated by using Gateway system (Invitrogen). The 2832bp DNA fragments which contains a cis-regulatory region of *mespb* gene were amplified from medaka (Taiwan local population medaka *Oryzias sinensis*) by using the specific primers;md-mesp-m2798F (5’-GTACATGGGACATCGCTGATTAGAGCAATAC-3’) and Mdmespb-p33R (5’-GTAGTCAAGAAAAGGGACCG

AGGA-3’) (see also Terasaki et al. (2006)). This fragment was inserted into the Gateway entry vector (pCR8/GW/Topo). The resulting vectors were used to transform *Escherichia coli* DH5α and sequenced. After determining the orientation of the insert, the insert fragment was inserted into the destination vector (*pT2G-GW-gate2p-ERT2Cre*) with the Gateway LR Clonase enzyme mix kit (Invitrogen).

### Zebrafish

Zebrafish handling was performed under standard conditions following institutional guidelines. We received ethical approval from Institutional Animal Care & Utilization Committee, Academia Sinica. Zebrafish were maintained and embryos were obtained and raised in the aquarium facility at Yilan Marine Research Station, Institute of Cellular and Organismic Biology, Academia Sinica. Lab strain wild-type zebrafish, Tg(*P067Olactb:loxP-dsR2-loxP-EGFP*) (Yoshinari et al., 2012) and Tg(*Olmespa-ERT2-Cre-ERT2*) were used in this study. Tg(*P067Olactb:loxP-dsR2-loxP-EGFP*) strain was received from Kawakami Atsushi’s lab and maintained in the aquarium facility. Tg(*Olmespa-ERT2-Cre-ERT2*) were established by *Tol2*-mediated transgenesis (Suster et al., 2009). By crossing *actbRG* and *mespCre*, we generated the hybrid transgenic line containing the above two transgenes (mespRG) (Fig. 1).

### Fluorescence imaging

The embryos are harvested from the mespRG parents. They incubated in the 5 cm Petri dish with E3 buffer at 24 to 28°C. At the bud to early somite stage, 4-Hydroxytamoxifen (4-OHT) was added to the Petri dish, and these embryos were incubated. From 3 to 4 days post fertilization (dpf), the hatched larvae were sorted based on the EGFP expression. The embryos expressing the EGFP signal at the post-cloacal level were isolated into the 5 cm Petri dish and observed under the fluorescent microscope Szx-16 (Olymups). The fluorescent patterns were recorded by photographing under the light and fluorescence view. The expression of EGFP was recorded every two to four days. During the photographing, the fish larvae were lightly anesthetized with MS222. The photographed larvae were maintained in a grass beaker (250 ml) until they developed caudal fin muscles.

### Histology and immunohistochemistry

The basic experimental procedure of conventional histology and immunohistochemistry is based on previous reports (Li et al., 2015, 2019). The zebrafish embryos and juveniles were soaked in Bouin’s fixative, embedded in paraffin, section to 5 *µ*m using a slicer (RM 2245, Leica). The sections were placed on the slide grass (Platinum-pro, Matsuinami). For the conventional histological analyses, the side was deparaf-finized with xylene, gradually hydrated with ethanol series, and stained with Alcian blue, hematoxylin, and eosin. For the section immunolabeling, the slides were incubated at 65°C, deparaffinized with xylene, and immersed in 100% methanol. The deparafinized slide was immersed in 1-3% hydrogen peroxide/methanol solution for more than 1 h, to remove endogenous peroxide. Thereafter, the slides were hydrated with ethanol series, blocked in 3% skim milk PBTx (1% Triton X-100 in PBS) for 1 h, and incubated with the primary antibody (the anti-green fluorescent protein rabbit IgG fraction [A11122, Invitrogen]) diluted in Superblock Blocking Buffer in TBS (#37535, Thermo Fisher Scientific) (1:100) at room temperature overnight. On the following day, the slides were washed with PBST three times for 3 min each and incubated with the secondary antibody (Goat pAb to Rb IgG [Abcam, ab6721]) diluted in the blocking buffer (1:1000) at room temperature three to six hours. After the secondary antibody incubation, the slides were washed three times each in PBST for 3 min each. The signals were detected by Thermo scientific Pierce Metal Enhanced DAB substrate Kit (#34065, Thermo Scientific). These slides were examined by the standard microscope (BX43, Olympus).

## RESULTS AND DISCUSSION

We succeeded in observing the EGFP distribution patterns in ten individuals from 4 days post-fertilization to more than 10 dpf (Supplementary Fig. S1). Presumably due to the genetic background and subtle differences in the strength and timing of the tamoxifen stimulation, these zebrafish specimens showed various patterns of the EGFP expression in different strengths of fluorescent light; these ten individuals are designated as 2018-1103-01, 2018-1130-01, 2018-1130-02, 2019-0617-01, 2019-0617-02, 2019-0617-04, 2019-0617-05, 2019-0617-03, 2019-0617-06, 2018-0629-06 (Supplementary Fig. S1). Of those, three of them, including, 2018-0629-06-, 2019-0617-03, and 2019-0617-06 show the signals which allow us to investigate how the caudal muscle tissues are developed from early larval stages (Fig. 2). In addition, we succeeded in conducting the histological analyses in 2019-0617-06 and 2018-0629-06, indicating that our genetic mosaic analysis works for the studies to investigate where the muscle tissue primordial cells appeared at the early larvae and differentiate into which caudal muscles (Fig. 3). By analyzing the three individuals more carefully, we clarified that a large part of the caudal muscle tissues derived from the pigmentless region at the ventral side of the caudal region (Fig. 2 and 3); the region showed condensation of mesenchymal cells which is indicated as the “gap in the melanophore stripe along the ventral myotomes” by Parichy and colleagues (2009).

**Fig 2.**
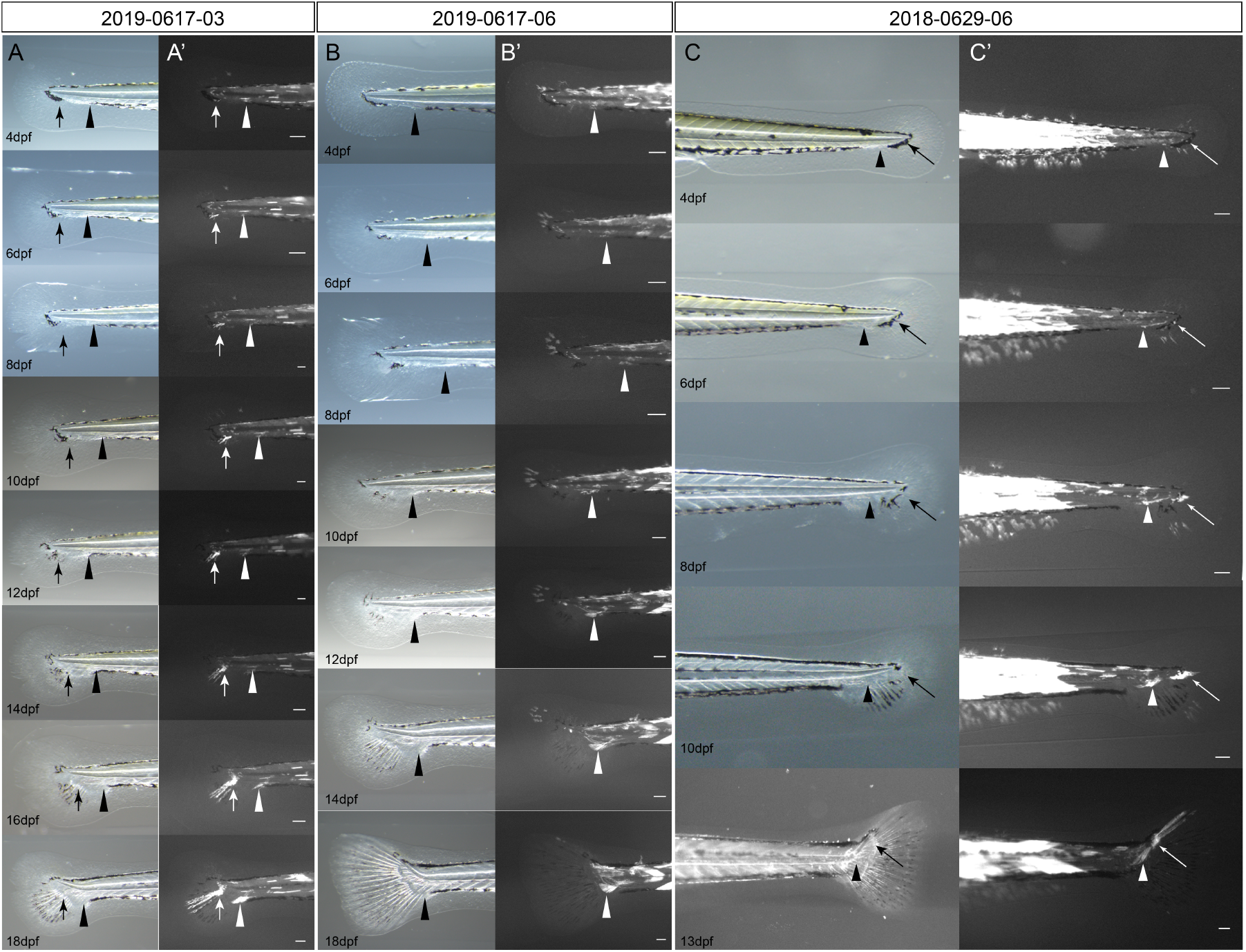
Tracing of the EGFP-positive cells at the caudal region. The lateral side views of the caudal region of the 2019-0617-03 (A), 2019-0617-06 (B), and 2018-0629-06 specimens. The day post-fertilization is labeled at the lower bottom corner of each panel. The arrowhead indicates the significant EGFP-positive cell populations. The arrows indicate the EGFP-positive cell populations differentiating into the fin rays. The notochord lengths (NTL) of the specimens are 3.478 mm for 2019-0617-03, 3.436 mm for 2019-0617-06, and 3.402 mm for 2018-0629-06.

**Fig 3.**
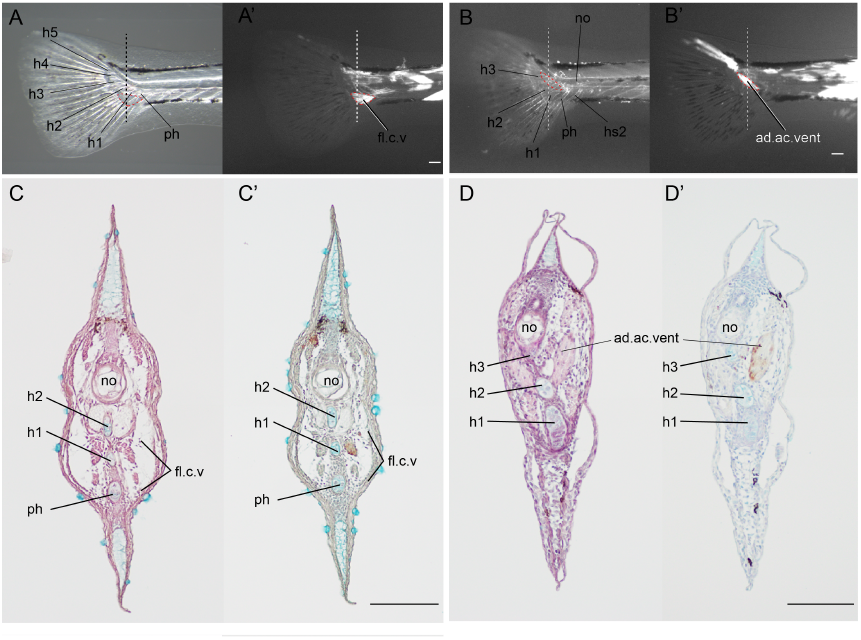
Histological analysis. The right-side view of the caudal region of the 2019-0617-06 (18 dpf) and 2018-0629-06 (13 dpf) (A, A’, C, C’). The transverse sections of caudal levels (B, B’, D, D’). Panels B and D show images of conventional staining. Panels of B’ and D’ indicate the immunohistochemistry of EGFP staining.

Moreover, from the results of 2019-0617-03 (3.478 mm in NTL), the primordial cells of the ventral caudal muscles are derived from the most anterior region of the mesenchymal cells condensation area (the arrowheads in Fig. 2A). Although there remains ambiguity as to whether the EGFP-positive cells differentiated into the deep layer muscle of the caudal muscle or not due to the absence of the results of the histological analyses, it seems that the muscle fibers belong to the flexor caudalis ventralis from the location and running orientation of the muscle fibers. In 2019-0617-06 (3.436 mm in NTL), several cells at the anterior part of the pigmentless region exhibit the EGFP and also differentiate into the flexor caudalis ventralis (the asterisks in Fig. 2B, 3A). In 2018-0629-06 (3.402 mm in NTL), the middle levels of the pigmentless region showed the EGFP-positive cells and finally, this cell population formed the adductor caudalis (Fig. 2C, 3B).

Our observations in the cell populations shown in Fig. 2BC correspond to those in the previous study (Siomava et al., 2018). In comparison, the zebrafish individual observed in the previous study was 4.4 mm in NTL, whereas the individual observed in our study was approximately 3.4 mm in notochord length. This difference indicates that our results represent an earlier developmental stage of the caudal region compared to the previous study (Fig. 2). More specifically, our study captures the status of muscle primordial cells before they differentiate into the long muscle fibers of the teleost-specific caudal fin muscles, as observed in the previous study (Siomava et al., 2018).

Although our study lacks quantitative data, it demonstrates that small cell populations eventually develop into a substantial portion of muscle fibers by the late larval stage (Fig. 3). These results are also consistent with previous observations of caudal region development in medaka, where a small number of muscle fibers ultimately develop into a significant part of the caudal muscle system (Ishikawa, 1990), suggesting a conserved developmental process shared by these species. Taken together, these findings support the idea that only a very small population of cells underwent modifications during the morphogenetic process over evolution in ray-finned fish, leading to the lineage-specific caudal muscle morphology observed in teleosts.

While the teleost-specific caudal fin skeletal morphologies are explained as the consequence of the fusion of several metamerically arranged skeletal elements (Bensimon-Brito et al., 2012; Schultze and Arratia, 1989), it is thought that more complex alterations occurred in the caudal muscles of teleost fish lineage. This complexity of the caudal muscle development suggests that it might be difficult to investigate its underlying developmental mechanisms. However, recent progress in the zebrafish studies (including the investigation of the relationship between *hox13* paralogues (Cumplido et al., 2024) and caudal fin morphology, and the high-resolution tracing research on the zebrafish muscle (Kumar et al., 2024) provided us an opportunity to apply these approaches to reveal how such complex muscle morphology and its developmental mechanism was established. We hope that our study contributes to understanding the evolution of this complex caudal musculature in fish.

## ACKNOWLEDGEMENTS

This work is supported by the NIG Scientist and National BioResource Project from the Ministry of Education, Culture, Sports, Science and Technology of Japan. We want to express our sincere gratitude to Dr. Atsushi Kawakami of Tokyo Institute of Technology for kindly providing the transgenic fish (*Olactb:loxP-dsR2-loxPEGFP*). We also thank the following current and ex-members of the Yilan Marine Research Station, Institute of Cellular and Organismic Biology; the late Hung-Tsai Lee, Chia-Chun Lee, Chihi-Chiang Lee, Tsai Han Chuan, and Chen Yi Wang for maintenance of aquarium systems; Chi-Fu Hung, Jhih-Hao Wei, Fei Chu Chen, and Ing-Jia Li for administrative support; Teng Yu-Feng (Big Wind Technology Co. LTD), Shi-Chieh Liu for designing aquarium systems, Jo-Hsin Chen for molecular cloning. We also thank the Taiwan Zebrafish Core Facility at Academia Sinica.

## FUNDING

The funding was provided by National Science and Technology Council (Grant No. 112-2311-B-001-033), Academia Sinica through the Postdoctoral Scholar Program (Grant No. 235g), JSPS KAKENHI (Grant No. JP22K06232), and Takeda Science Foundation (Grant No. 2022036015).

## AUTHOR CONTRIBUTIONS

**Conceptualization**: Kinya G. Ota 太田欽也 (KGO), Gembu Abe 阿部玄武 (GA). **Funding Acquisition**: KGO, GA. **Investigation**: KGO, GA.. **Methodology**: KGO, GA. **Project Administration**: KGO. **Resources**: KGO, GA. **Supervision**: GA. **Image acquisition**: KGO. **Writing – Original Draft**: GA. **Writing – Review & Editing**: KGO, GA.

## COMPETING FINANCIAL INTERESTS

The authors declare no competing financial interests.

## Supplementary Information

**Supplementary Fig S1.**
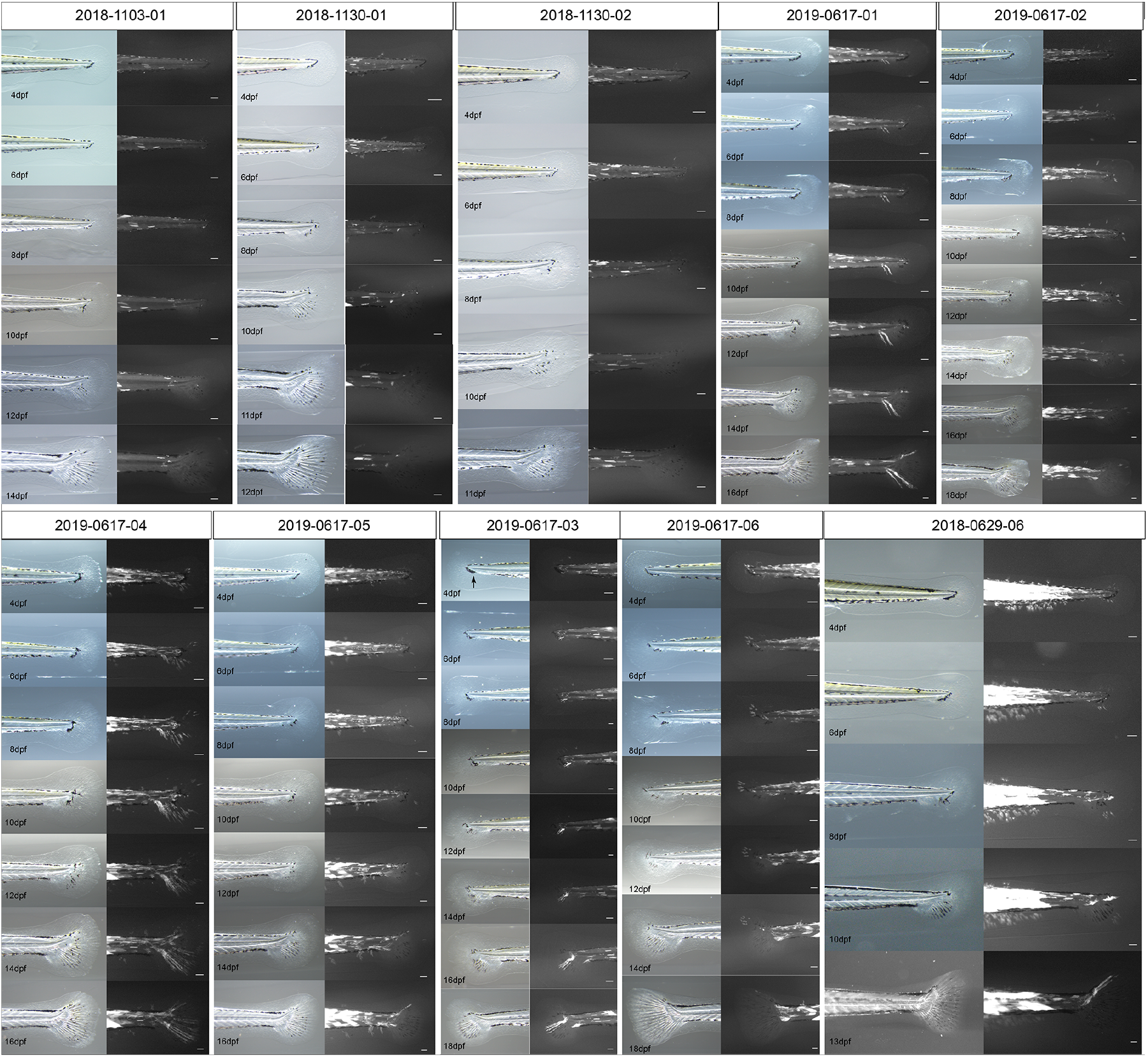
Caudal lateral views of mespRG zebrafish individuals. The days post-fertilization is shown in the left-lower corner. Panels on odd and even number columns display the light and dark views.

## Notes

### Competing Interest Statement

The authors have declared no competing interest.

